# Long read sequencing reveals changes in measured isoform structure and abundance due to RNA processing

**DOI:** 10.64898/2026.11.03.764063

**Authors:** Rotem Aharon, Alicia Oshlack

**Affiliations:** Peter MacCallum Cancer Centre, Parkville, Australia; Sir Peter MacCallum Department of Oncology, University of Melbourne, Parkville, Australia; School of Mathematics and Statistics, University of Melbourne, Parkville, Australia

**Keywords:** Long read RNA sequencing, Transcriptomics, PCR amplification bias, Reverse Transcription Errors, RNA m^6^A modification, RNA methylation, Differential Transcript Usage, Isoform Switching, Alternative Splicing

## Abstract

Oxford Nanopore Technology enables direct RNA sequencing without reverse transcription (RT) and cDNA sequencing without PCR amplification, allowing investigation of how RNA preparation protocols bias sequencing data. PCR amplification is known to introduce biases related to expression levels, transcript lengths, and GC content. However, biases introduced by RT have been described more recently and their quantification remains more elusive. Using seven cell line datasets from SG-NEx, we compared nanopore sequencing of direct RNA, direct cDNA, and PCR-amplified cDNA. We identified hundreds of differentially expressed genes and isoforms between protocols, indicating significant changes in observed expression. Differentially expressed isoforms displayed expected amplification biases, and further we observed decreased PCR counts associated with higher m^6^A methylation levels on native RNA. We also found differential transcript usage in hundreds of genes across cell lines, showing substantial changes in measured isoform proportions between protocols. Analysis of structural differences between isoforms that switch dominance revealed that while some events involve simple RNA fragmentation, most involve complex structural differences including novel junctions (exon skipping, intron retention, etc’). Compared to non-DTU genes, inserted introns (exitrons) were significantly more common after RT and amplification. Furthermore, visualisation of raw coverage data proved crucial for qualitative validation of experimental findings.

## Introduction

Long-read RNA sequencing (lrRNA-seq) technologies enable higher confidence and accuracy in the identification of the alternative transcriptional isoforms that make up the transcriptome. This has resulted in increasingly detailed descriptions of transcriptomes across different cell and tissue types, disease states, developmental stages, and cancer (1–4).

While traditional short read RNA sequencing (srRNA-seq) technologies require preparation steps including reverse transcription (RT) and PCR amplification prior to sequencing, lrRNA-seq protocols now allow sequencing of the RNA directly without RT (direct RNA), or the sequencing of cDNA without PCR amplification (direct cDNA), along with the more common RT and PCR preparation protocol (PCR cDNA).

PCR amplification is known to induce biases associated with expression levels, transcript lengths and GC content (5–7), and over the years several computational methods were developed to account for these (8–13). Biases introduced by the RT step have also been identified. These include the reproducible identification of false exitrons or “gaps” in exons (14), and the report of higher levels of chimeric reads in cDNA (either from direct cDNA or PCR cDNA) as compared with direct RNA sequencing (15). A comprehensive list of possible artefacts induced by RT and PCR has been reported in (16). Characterisation of RNA degradation rate in direct RNA samples is available (17), and a computational method has been suggested to mitigate biases caused due to the variation in the distribution of RNA fragment length between protocols (18).

Since the introduction of lrRNA-seq technologies, several computational tools have been developed specifically for the analysis of this type of data. Isoform annotation and quantification methods include Bambu, IsoQuant, FLAIR, FLAMES, TALON, Mandalorion, IsoTools and OarFish (19–28). Further, SQANTI3 was developed to enable quality control and structural annotation of isoforms detected from lrRNA-seq data (29–31). Downstream analysis steps such as differential expression and differential transcript usage analysis are mainly performed using methods that were initially developed for short read analysis; their performance on lrRNA-seq data was benchmarked in (32). However, more recent improvements to downstream analysis include adjustments designed to accommodate for the specific characteristics of lrRNA-seq counts, such as quantification of uncertainty and read-to-transcript ambiguity (33–35). A comprehensive review of the lrRNA-seq analysis workflow is presented in (36).

Benchmarking efforts designed to compare sequencing methods, library preparation protocols, and analysis tools for long-read RNA sequencing data highlight the immense improvement both technologies and analysis tools have undergone in recent years (32,37–41). Despite this, some challenges still remain, mainly around the accurate identification of transcripts starts, ends, and splice junctions, and the allocation of truncated sequence reads to the correct transcript isoform (36,42,43). Different long-read sequencing technologies and protocols generally display good agreement between gene level counts, but results are more variable at the transcript level (39,41).

In this study, we aim to assess how different aspects of RNA preparation affect lrRNA-seq data. Specifically, we analyse public data provided in SG-Nex (39), where RNA collected from 7 different cell lines was sequenced using three ONT preparation protocols (direct RNA, direct cDNA, PCR cDNA), short-read Illumina sequencing with replicates, and a single replicate of Pacbio iso-seq. Using this data, we evaluate the changes in gene and transcript expression that are induced by the protocol, and look at potential associations with transcript length and GC content. Further, we investigate the potential role of RNA epigenetic modifications in influencing the bias between protocols. Next, we evaluate differences in transcript usage between protocols and find evidence of several genes where the dominant isoform is different between the different protocols (isoform switching). Interestingly, many of these switches in dominant isoforms have evidence of internal splicing differences rather than simple changes to the start or end of transcription. We highlight a second, much smaller group of differential transcript usage (DTU) genes, in which a ‘novel’ transcript is detected in the PCRcDNA samples but is not observed in the dRNA. These cases are rare but indicate that the introduction of ‘new’ alternative isoforms not observed in the dRNA can occur during the RT or PCR amplification stages. Last, we discuss the interplay between differences that can be observed in read coverage data, and those which seem to be a result of errors in allocations of reads to transcripts, and suggest that visualisation of the read-coverage data provides qualitative validation of findings obtained based on transcripts counts and annotation alone.

## Methods

Below we describe the methods and tools used for this analysis. Detailed code for the analysis and figures is available at https://zenodo.org/records/21132314.

### Data overview (Library size, dimensional reduction and sample inclusion)

The data analysed here is publicly available from the SG-Nex database (39). The SG-NEx data was accessed in June 2025 at https://registry.opendata.aws/sg-nex-data.

We started by downloading 107 .bam files (samples), which include read alignment to the human genome (hg38) for each of the replicates in each of the cell lines. Read alignment to the genome was done by the curators of the SG-Nex dataset using minimap2 v.2.22 with parameters --ax splice --junc-bed, using the junction bed file to correct splicing junctions, and the additional parameters --k14 and --uf were used as for direct RNA sample as recommended. Base calling was performed using Guppy v.6.4.2. For further details on methods used in the curation of this dataset see (39).

bambu v.3.8.3 (19) was used to identify and quantify the genes and transcripts in all of the samples together (NDR=0.1). This resulted in a SummarisedExperiment R object, which includes transcript counts and full length read counts as separate assays. Gene counts were obtained by using bambu::transcriptToGeneExpression, which effectively adds the transcript counts for each gene.

For each sample, library size was obtained by aggregating transcript counts. Samples with very low or very high library size <400,000 or >40,000,000 were excluded, as well as those that had full length to all transcript ratio lower than 0.2.

This left us with 72 ONT samples and 6 PacBio samples. Supplementary Table S1 summarises sample inclusion in each of the analysis sections.

MDS dimensional reduction plots were generated using edgeR:: plotMDS v.4.4.2 (33,44) for each of the assays independently, with increasing top=2000, to account for relatively high correlation between transcript counts.

Since we decided to proceed with the analysis to compare only between ONT lr-RNAseq protocols, we run bambu again to obtain more precise annotation and quantification of transcripts based only on ONT samples. The rest of the analysis presented below is performed on the results of this later ‘bambu’ quantification run (updated library size and MDS plots based on this quantification are presented in Supplementary Fig S1). After applying the same inclusion criteria we were left with 72 ONT samples across the three protocols.

### Differential Expression Tests

Differential Expression analysis was performed separately for each cell line, with a design matrix constructed based on the sequencing protocol. For each of the seven cell lines, three count matrices were obtained from the relevant samples in the SummarisedExperiment: gene counts, transcript counts, and transcript counts obtained based on full length read only. Using limma (voom) v.3.62.2 (45), raw reads counts were filtered to remove low expression, with min_count = 20 and min_total_count = 50 for gene counts, and min_count = 10, min_total_count = 20 for transcript counts. After filtering, voomWithQualityWeights was utilised to adjust for varying sample quality, and a linear model was fitted with lmFit. Then limma::decideTest was used with p.value=0.05, with lfc thresholds of 1.0 for genes and 0.5 for transcripts, and topTables were obtained for each of the comparisons in each of the cell lines.

### Sequence length and GC content

Based on bambu extended annotation output and the relevant reference genome, Bedtools *nuc* (46) was used to obtain the number of occurrences of each nucleic acid in each of the annotated transcripts. Sequence length is the sum of all bases, and GC content is the ratio of sums of G’s and C’s to the total length.

### m^6^A methylation

m^6^A detection results are available from the SG-Nex database for a subset of the direct RNA samples in the core dataset (A549, K562, Hct116, HepG2, MCF7). For each of the cell lines, m^6^A site modification probability was calculated using m6Anet *infer* (47) based on the 2-3 replicates that were provided for each of the cell lines. For each isoform, we first counted the number of sites that were methylated with probability >0.9 (the threshold for true modification suggested by the authors), and then calculated the fraction of these sites out of all possible methylation sites on that isoform, to provide a single value that represents methylation fraction for each isoform.

### DTU tests

Testing for significant DTU between the PCR cDNA and direct RNA samples was performed separately for each cell line using DRIMSeq v.1.34.0 (48), based on all transcript counts. False discovery rate (FDR) was controlled using a Two-stage test implemented in stageR v.1.28.0 (49), as recommended by the authors in (50).

More specifically, parameter values used default settings in DRIMSeq:dmFilter( ) except: min_samps_feature_expr = 2, min_feature_expr = 5, min_samps_gene_expr = 2, and min_gene_expr = 10.

### Isoform switching genes

In order to identify isoform switching genes, for each significant DTU gene, all isoforms were ordered by their proportions in each condition. If the most dominant transcripts are different between the different protocols, this is defined to be an isoform switching gene. The most dominant transcript in direct RNA is set as ‘reference’, and the one most dominant in PCR cDNA is set as ‘test’.

### Background genes

For non-significant DTU genes, the isoforms were ordered by their proportion in the direct RNA. The most dominant isoform was set as ‘reference’ and the second most dominant was set as ‘test’.

### Sampled sets of background genes

Since the mean expression of the DTU genes was generally higher than that of the background genes, for each cell line we randomly sampled 5 sets of 100 background genes with an expression distribution consistent with that of the isoform switching genes from the same cell line.

### Novel isoforms in PCRcDNA

A novel transcript in the PCRcDNA group is defined to be a DTU transcript for which the proportion in the PCRcDNA groups is at least 0.01, and effectively zero (<0.001) in the direct RNA group. The most dominant transcript in direct RNA is set as ‘reference’, and the novel transcript present only in the PCR cDNA samples is set as ‘test’.

### SQANTI3 labels

SQANTI3 v.5.2.1 (29) was used to obtain the SQANTI3 structural category of the test transcript with respect to the reference transcript. For each cell-line, the .gtf file provided as reference to SQANTI3 included only the isoforms considered as reference in that cell line, and the query file included only the isoforms considered as test in that cell line. Because sqanti3 removes isoforms that it considered too similar from the reference prior to labelling, and since it pairs each test isoform back to the reference isoform it considered as the most similar to it from the list of reference isoforms, not all reference vs test pairs we are interested in can be labelled by sqanti3. Figure 3f therefore presents only pairs for which SQANTI3 labels are available.

### Identify and characterise structural differences

Based on all exon start and end positions on the genome, all unmatched positions were identified and marked on the test transcript with respect to the reference transcript. This identification step accommodates for small variations in transcript annotation that could be a result of an inaccurate identification of the splice junction, by allowing for unmatched positions that are in very close proximity (here bp=20, or +/-10bp around exon start and end positions) to be considered as matched. Unmatched positions were characterised and labelled based on their relationship to the surrounding features, for example: ‘exon skipping’, ‘intron insertion’, etc. This can result in multiple events allocated to any specific comparison between transcripts. When these events occur consecutively (without any matched exon start or end position between them) they are considered linked events. For example, consecutive skipped exons are labelled ‘linked deletion’, and consecutive inserted exons are considered ‘linked insertion’, as it is likely they are the result of a single splicing event. An alternative exon, which in our case is identified as an exon skipping event followed by an exon insertion, or vice versa, is labelled a ‘linked combination’ event (similar to an ‘exon extension’ event followed by an ‘exon skipping’, or many other possible combinations). Additional details regarding structural differences detected in the test compared with the reference are also collected, such as their position, length, and a label to indicate the position on the transcript (3’, 5’, or internal).

We also quantify and visualise the structural differences detected, along with other features which enable integration with coverage data (e.g. for qualitative validation) and with annotation level information (such as CDS and ORF).

## Results

### Data overview

Here we aim to perform a rigorous comparison between long-read RNA sequencing protocols (lrRNA-seq), in order to quantify and characterise biases present between them. For this purpose, we chose to analyse data from the SG-NEx dataset (39), which includes lrRNA-seq data obtained using different sequencing technologies and different preparation protocols with multiple replicates, for multiple cell lines.

We start by downloading the .bam files containing all reads aligned to the human genome, for all cell lines sequenced using at least three out of the four lrRNA-seq protocols (PacBio IsoSeq, and three ONT protocols: dRNA, dcDNA and PCRcDNA). These filters leave 101 ONT samples and 6 PacBio samples across 7 cell lines (A549, H9, Hct116, HepG2, HEYA8, K562 and MCF7, where H9 had no PacBio data available). There is large variation in the number of reads in each sample (Fig. 1a). We quantify genes and transcripts using bambu (19), and exclude samples with very low or very high library size <400,000 or >40,000,000 reads (Fig. 1a). Bambu can assign reads as “full-length” based on read class characteristics such as read coverage, read alignment and read sequence characteristics. These reads are thought to more fully capture the structure of the transcripts without truncation (39). Fig. 1b shows the ratio of the number of full-length reads to total reads as a function of the total library size of all transcript counts. We note that for most samples, apart from PacBio, the fraction of full-length transcripts is less than 0.6 of the total library size. To ensure data quality we further exclude samples with a very low fraction of full-length transcripts (<0.2).

**Figure 1.**
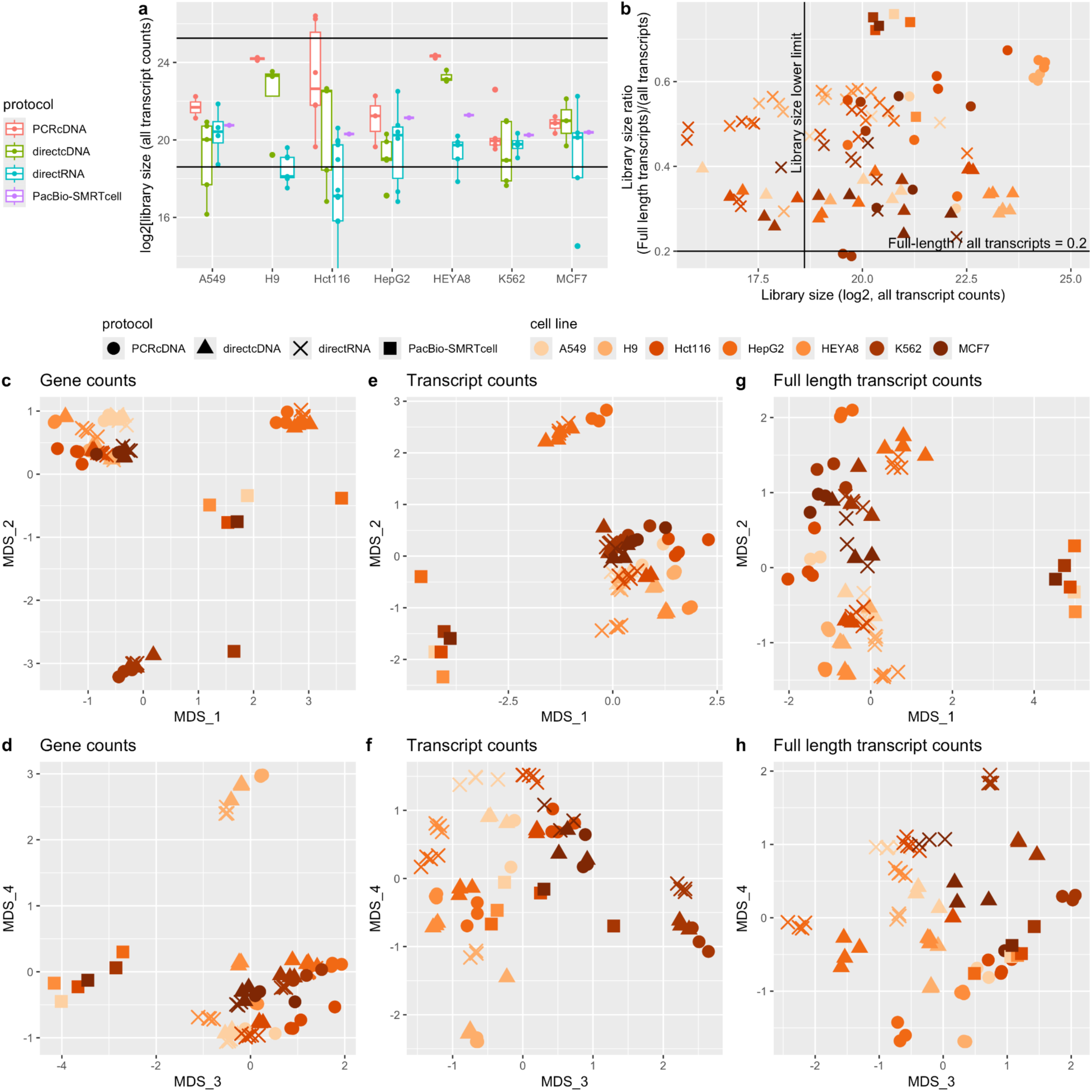
(Data overview): a) Library size (number of mapped reads) in each sample, across preparation protocols and cell lines. The samples between the horizontal lines were retained for analysis. b) Fraction of full length transcript reads as a function of log2(library size). MDS plots of gene counts (c,d), total transcript counts (e,f) and full length transcript counts (g,h). Samples separate based on cell line (colour) but also on preparation protocol (shape) especially at the isoform level.

Multidimensional Scaling (MDS) dimension reduction plots of the 79 remaining samples reveal several interesting features of variation between samples. At the gene expression level, samples clearly separate by cell line, except for the PacBio samples, which cluster separately from ONT (colours, Fig. 1c-d). At the transcript expression level, PacBio also clearly shows differences in overall quantification of samples. This has been shown and discussed in previous studies (39,41).

The largest source of variation between the ONT samples is due to the cell line (Fig. 1c-e). However, at higher dimensions, samples further separate based on preparation protocol, particularly in transcript expression from dimensions 3 and 4 (Fig. 1f). When looking at full length transcript counts, there is evidence of the protocol contributing to variation even in the first dimension (Fig. 1g). These observations together give strong evidence that the protocol influences the measured expression level of transcripts. Because we seek to understand biases caused by RNA library preparation, and because the PacBio samples do not have replicates for statistical analysis between protocols, in the following section we study the differences between only the ONT data obtained using different library preparation protocols. Supplementary Fig. S1a-h reproduces Fig. 1 with the PacBio data excluded from as early as the transcript quantification step performed with ‘bambu’.

### Differences in expression levels between preparation protocols

We perform differential expression analysis between the three different ONT preparation protocols. Specifically, for each cell line, we test for Differential Gene and Differential Transcript Expression (DGE and DTE respectively), using limma-voom (45,51), over three contrasts: direct RNA vs. direct cDNA, direct RNA vs. PCR cDNA and direct cDNA vs. PCR cDNA.

The numbers of significant features detected in each cell line are presented in Fig. 2a and 2b for the DGE and DTE tests respectively. Across each of the seven cell lines we detect hundreds to thousands of significantly differentially expressed genes and isoforms. The cell lines with the largest number of differentially expressed features are those that have larger average difference in library sizes between protocols (H9, HEYA8 and Hct116). This may be a result of technical artifacts that affect the quality of sequencing reads, or analysis artifacts in the assignment of reads, or statistical analysis, which has higher power to detect significant differential expression with higher expression levels. Significant DE features are biased towards higher expression levels as expected (Supplementary Fig. S2a).

**Figure 2.**
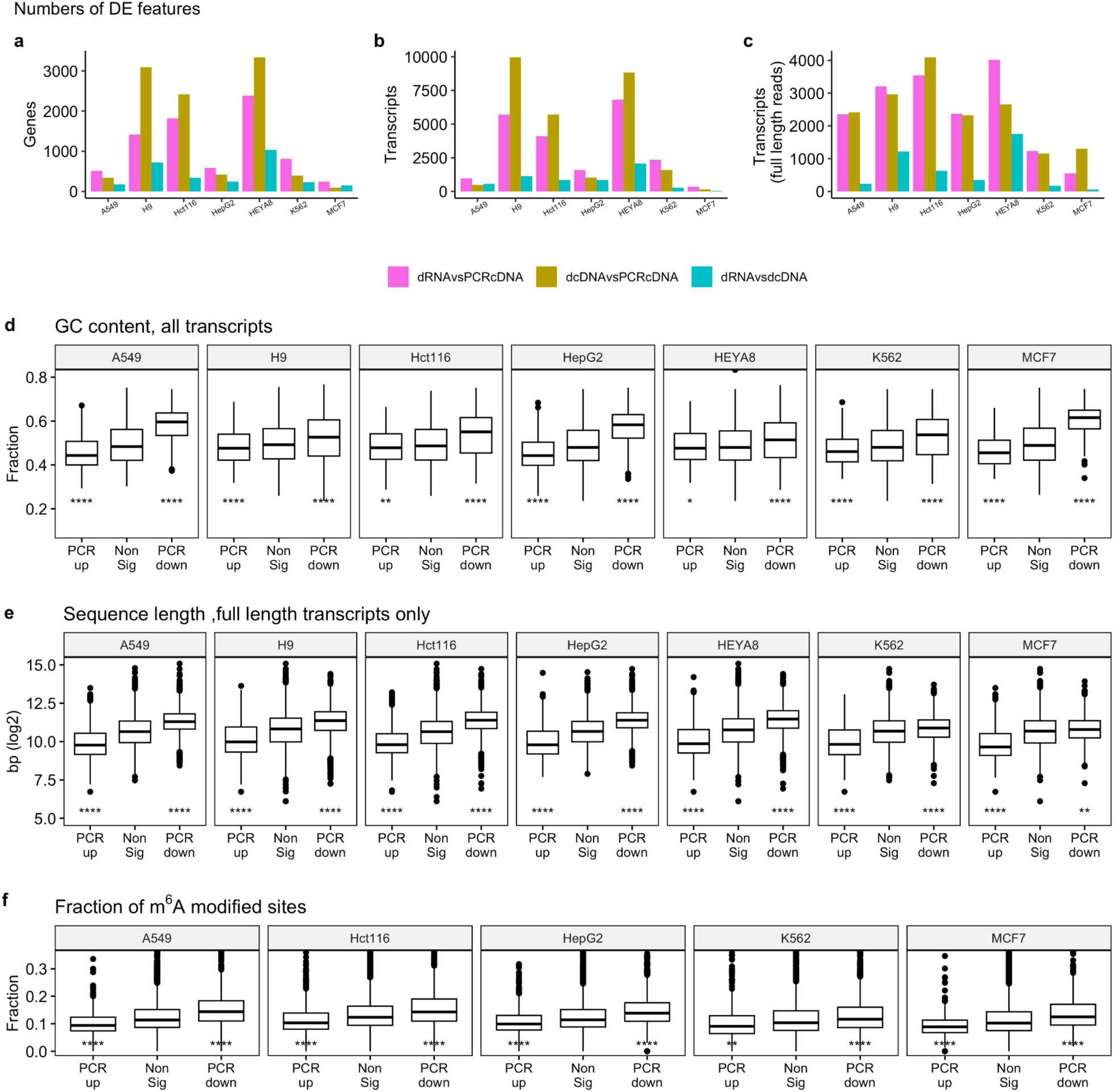
Differential Expression between sequencing protocols: a) Number of significant DE genes found for each cell line in each of the contrasts tested ( adjP=0.05, lfc=1). b) Number of significant DE transcripts found for each cell line in each of the contrasts tested ( adjP=0.05, lfc=0.5). c) Similar to b) but calculated using full length transcript counts only. d) GC content of all transcripts split by DE label obtained when comparing PCR cDNA samples with direct RNA (with testing performed on all transcript counts). e). Similar to c) but plotting the sequence length (DE testing performed on full-length transcript counts). f) Similar to c) but plotting the fraction of methylated sites in the transcript (DE testing performed on all transcript counts, plotting only transcripts for which methylation data was available in SG-Nex). P-values are obtained using a two-tailed t.test with respect to the non DE group. ns: p > 0.05, *: p <= 0.05, **: p <= 0.01, ***: p <= 0.001, and ****: p <= 0.0001

**Figure 3:**
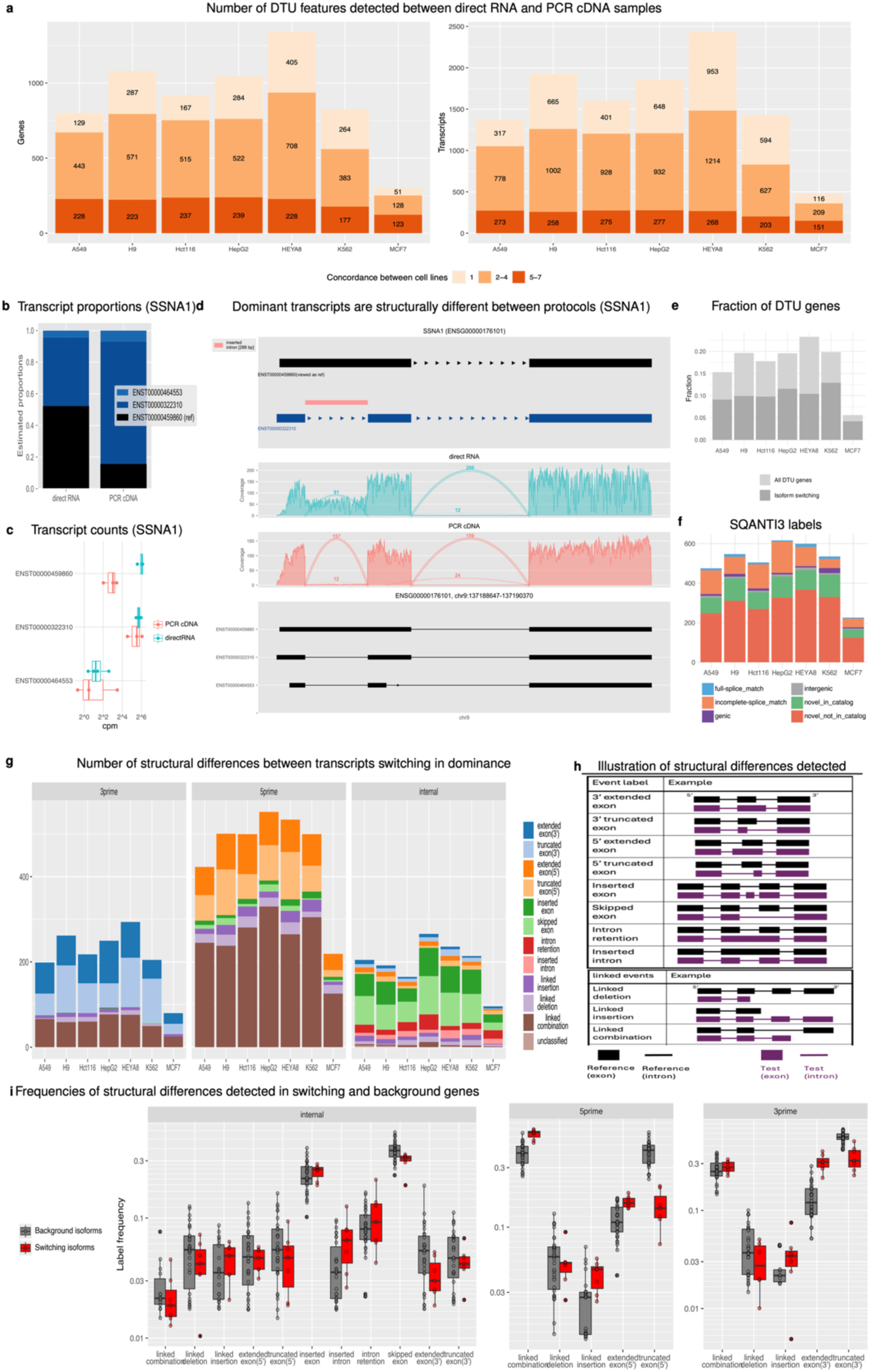
Differences in the dominant isoform between protocols a) Number of DTU genes and transcripts detected between direct RNA and PCR cDNA samples. Different colours indicate concordance between cell-lines. Light orange is the number of features that were DTU only in that cell line, and darker orange is the number of features detected in 5 or more of the cell-lines. b-d) Example of a DTU gene (SSNA1) where the dominant transcripts are different between protocols. b) Estimated proportions in direct RNA (left) and PCR cDNA (right), estimated using DRIMSeq from ‘bambu’ raw transcript counts for the same gene. c) Transcript counts (log2cpm) obtained by ‘bambu’ for this gene. d) Switching isoforms in SSNA1 compared: Dominant transcript in direct RNA is considered reference in black (ENST00000459860), dominant transcript in PCR cDNA is considered test in dark blue (ENST00000322310). In this case, the reference transcript is clearly longer than the test one. Structural differences between the two transcripts were identified, characterised, and then labelled on the test transcript with respect to the reference. Here an intron insertion was identified. While this difference was initially detected based on annotation and transcript quantification alone, coverage plots from representative samples provide qualitative validation of this based on aligned sequence data. Annotation plot of all transcripts detected in these samples by ‘bambu’ is provided for completeness. e) Fraction of DTU genes detected across cell lines, with the fraction of genes with dominant switching in darker grey. f) SQANTI3 labels obtained for each of the switching isoform pairs in each cell line. g) The number of structural differences detected between switching isoforms in each of the cell lines. h) Schematic representation of structural differences detected between reference (black) and test (purple) transcripts. i) Fraction of structural differences detected between reference and test transcripts in switching isoform genes (red), compared to the fraction of differences detected in sampled sets of background genes (grey).

When comparing protocols, the highest fractions of DE genes and transcripts are found when comparing direct cDNA to PCR cDNA, but the direct RNA vs PCR cDNA comparison also displayed many changes. We find the lowest number of differential features when comparing direct RNA to direct cDNA, suggesting that many expression changes are associated with the PCR amplification step. However, PCR cDNA samples also have much higher library sizes in most cell-lines, and therefore would lend more power to detecting significant DE. Since the most common and robust approaches to sequencing the transcriptome are direct RNA and PCR cDNA, we focus on investigating the biases specifically in this comparison.

PCR amplification bias is known to be associated with GC content, where sequences with very high or very low GC content do not get amplified as efficiently. Further, preferential amplification of shorter sequences has been previously reported (6,7). To investigate the effect of GC bias between protocols, we compute the GC content of all transcripts, and stratify transcripts based on the significance and direction of their differential expression between the PCR cDNA and direct RNA samples for each cell line. Fig. 2d shows that transcripts that are down-regulated in PCR cDNA compared with direct RNA have higher GC content on average, while those that are up-regulated after RT and PCR have significantly lower GC content across all 7 cell lines.

Next, we look at the sequence length of the transcripts. To try to avoid shorter reads caused by fragmentation of transcripts, we show only results based on DTE testing of full-length read counts. Fig. 2e shows that in all cell lines, the transcripts that are up-regulated in PCR cDNA compared with direct RNA have significantly shorter sequence length, while those down-regulated are longer. This indicates that, as expected, shorter transcripts are preferentially amplified compared with longer ones.

Direct RNA sequencing allows for the detection of epigenetic modifications of the RNA. We utilise this to investigate whether the presence of m^6^A methylation on RNA has an impact on the detected expression level of transcripts after reverse transcription and PCR amplification. m^6^A detection results are available from the SG-Nex database for a subset of the direct RNA samples. For each of the cell lines in the core dataset (which includes A549, K562, Hct116, HepG2, MCF7), we calculate m^6^A site modification probability using m6Anet *infer* (47) on the 2-3 replicates that were provided for each of the cell lines. Based on these probabilities and the threshold suggested in (47) we calculate the fraction of methylated sites for each isoform (see methods). Fig. 2f shows the fraction of methylation for transcripts identified as having differential expression between the direct RNA and PCR cDNA protocols for each cell line. Intriguingly, we find that the transcripts downregulated after RT+PCR have a significantly higher fraction of RNA methylated sites, and further, in most cases the transcripts with higher expression after RT and PCR have a lower fraction of m^6^A methylation. A possible explanation for this observation is that the presence of m^6^A on RNA reduces the efficiency of reverse transcriptions.

A cross-correlation plot of observed transcript expression, GC content, transcript length and m^6^A methylation (Supplementary Fig. S2b) shows as expected that the number of m^6^A modified sites increases with transcript length, and decreases with GC content. However, the fraction of m^6^A modified sites (which is the measure of m^6^A modification plotted in Fig. 2f) is not correlated with sequence length (r=0.04), and has positive correlation with GC content (r = 0.38). Further, the average expression fraction shows negative correlation with the fraction of modified sites (r = -0.17), but is not correlated with the number of modified sites (r=0.01).

Transcript length and GC content are inherent attributes of isoforms which do not change between cell lines. Therefore, we expect some consistency in the DE genes and transcripts detected between protocols across the cell lines, even though overall expression levels can be vastly different. This is in fact the case with nearly a third of all DE genes shared across the majority (>5/7) of cell lines (Supplementary Fig. S2c). Relatively high concordance was found also in DE transcripts across protocol comparisons (Supplementary Fig. S2d).

Overall, these findings highlight a complex interplay between the observed expression of transcripts across different sequencing protocols and the three features of GC content, transcript length and m^6^A methylation. Although the impact of these cannot always be separated, the presence of biases in data generated from different protocols should be considered when designing experiments.

### Significant differences in transcript usage (DTU) detected between protocols

While the expression of individual transcripts are correlated within genes, there are some cases where the dominant transcript within a gene changes between tissues or samples. Such events can be detected by testing for changes in the proportion of transcript expression within genes between conditions, referred to as differential transcript usage (DTU).

We perform tests for DTU between direct RNA and PCR cDNA samples for each cell line, based on all transcript counts using DRIMSeq (48,50). DRIMSeq analysis includes preliminary filtering of lowly expressed genes and transcripts, estimating the relative proportion of the transcripts within each gene, then testing for significant differences in these proportions between conditions. We use stageR (49) to control the false discovery rate (FDR = 0.05). Hundreds of DTU genes and transcripts are detected in each of the cell lines (Fig. 3a), with around 200 DTU genes and 300 DTU transcripts appearing in 5 or more of the cell lines, which we consider very high concordance. As in DE tests, genes that have significant DTU tend to have higher expression, likely because the test has more power (Supplementary Fig. S3b). Despite these similarities to the DTE results, testing for DTU is inherently different. In DTU, the transcripts within each gene are not assumed to be independent of each other, but rather the proportion of all transcripts in each gene are estimated together, and the DTU test returns results at the gene and transcript levels. Within a DTU gene, there can be 1 or more DTU transcripts, and their relative proportions can vary significantly in different directions between conditions.

### Investigating the differences between dominant switching isoforms

In order to explore the factors that drive the differences in relative transcript expression we observe between direct RNA and PCR cDNA samples, we first filter the list of all significant DTU genes and transcripts to a subset in which we identify a clear pattern of change in relative proportion. Specifically, we define an *isoform switching gene* to be a significant DTU gene in which the dominant transcript (ie - most highly expressed transcript) is different between the protocols. For example, SSNA1 is an isoform switching gene detected between the direct RNA and PCR cDNA samples of the A549 cell line (Fig. 3a-c). In direct RNA samples, counts are almost equally split between the top 2 transcripts (ENST00000459860 and ENST00000322310 contribute about 0.5 and 0.45 respectively), while in PCR cDNA samples, ENST00000322310 contributes nearly 0.75 of the overall expression.

From the total of 6177 significantly DTU genes identified across the 7 cell lines, 2103 (∼34%) meet our criteria for isoform switching (Fig. 3d). For each isoform switching gene we compare the two transcripts that are dominant in the different conditions. We define the dominant transcript in the direct RNA samples as the ‘reference’ (e.g. Fig. 3c top transcript, in black) and the dominant transcript in the PCR cDNA samples as the ‘test’ (Fig. 3c second transcript, in dark blue). For contrast, we select genes with non-significant DTU, which we refer to as background genes, with the most dominant transcript in dRNA labelled ‘reference’ and the second most dominant transcript as the ‘test’.

First, we check if mean gene expression, GC content and sequence length are associated with the change in relative transcript expression between protocols, like in the differential expression sections above. Gene level expression of the isoform switching genes is significantly higher than the mean expression of the background genes in all the cell lines (Supplementary Fig. S3a). This is expected since higher counts increase the power to detect DTU. Since GC content and sequence length are highly correlated between transcripts of the same gene, for each gene we calculate the differences in GC content and sequence length between the reference and test transcripts (i.e. GC_diff= GC_ref-GC_test). These differences in GC content and sequence length generally display different distributions between the DTU and background genes. More specifically, the differences in GC content are centred around 0 for the background genes and are slightly higher in the switching genes (Supplementary Fig. S3b), whereas the differences in sequence length are centred around zero for the switching genes and are slightly higher in the background genes (Supplementary Fig. S3c). To further investigate differences in sequence length, we also look at the sign of the difference in sequence length. In the background genes, the dominant transcript has a longer sequence in around 75% of the pairs, whereas in the switching isoforms, the dominant transcript in direct RNA has a longer sequence in about 50% of the genes (Supplementary Fig. S3e). Since the evidence for such association is not as clear nor conclusive as observed in the DTE results, we proceed to take a more detailed look into the structural differences between the switching transcripts.

We obtain SQANTI3 structural category labels for all isoform switching genes by categorising the test isoform (dominant in PCR cDNA) with respect to the reference (isoform dominant in direct RNA), shown in Fig. 3e. Interestingly, SQANTI3 identifies a large number of the test isoforms (dominant in PCR cDNA) as Novel In Catalogue (NIC) or Novel Not In Catalogue (NNC) with respect to their counterpart reference isoform (dominant in the direct RNA). A smaller fraction of these are characterised as Full Splice Matches (FSM) or Incomplete Splice Matches (ISM).

To provide a more granular description of the differences between the switching isoforms, we develop an approach to identify and characterise the differences in intron-exon structure between two transcripts of the same gene. Based on all exon start and end positions on the genome, all unmatched positions are identified and marked on the test transcript with respect to the reference transcript. Unmatched start and end positions are characterised and labelled based on their relationship to the surrounding features, for example: ‘exon skipping’, ‘intron insertion’, etc.

Multiple events can be allocated to any specific pair of transcripts. When these events occur consecutively, without any matched junctions between them, we consider them as linked events since they are likely the result of a single splicing event. Consecutively skipped exons are labelled ‘linked deletion’, and consecutive inserted exons are considered ‘linked insertion’. An alternative exon, which in our case is identified as an exon skipping event followed by an exon insertion, or vice versa, is labelled a ‘linked combination’ event (similar to an ‘exon extension’ event followed by an ‘exon skipping’, or many other possible combinations). Additional details regarding events detected in the test transcript compared with the reference are also collected, such as their position, total length, and relative position within the transcript (3’, 5’, or internal).

In the SSNA1 gene for example (Fig. 3a-c), we identify and characterise all structural differences between transcript ENST00000459860 (black, dominant in the “reference” direct RNA samples) and transcript ENST00000322310 (dark blue, dominant in the “test” PCR cDNA samples). We annotate an inserted intron in the test transcript with respect to the reference (Fig. 3c). The coverage plots (Fig. 3d) provide a qualitative validation to this observation which is initially based on testing transcript counts and annotations alone. It is evident that in the direct RNA samples a much higher proportion of reads cover the area labelled “intron insertion”, while in the cDNA PCR samples a higher proportion of the reads span the junction associated with this event. It is important to note that, in some cases, coverage plots of the relevant region do not validate the observations obtained based on transcript proportions and annotations alone (Supplementary Fig. S4). Such cases are likely due to misallocation of reads to transcripts during earlier stages of the analysis. Such misallocation can adversely affect the accuracy of transcript counts and annotations (38), and consequently distort the estimation of relative proportions on which DTU tests rely.

With this approach we characterise the differences detected in the test transcript with respect to the reference for all of the isoform switching genes. The overall number of structural differences detected in each cell line is presented in Fig. 3f, alongside a schematic description of all event types detected (Fig. 3g). At the 3’ and 5’ ends, tens to hundreds of extended or truncated exon events were detected. These events correspond to alternative start and end sites detected between the different protocols, labelled as FSM by SQANTI3. Many more skipped exons and linked deletions are detected on the 5’ end than on the 3’end. These correspond with the ISM label, and indicate that one or more exons are missing in the PCR cDNA dominant transcript with respect to the transcript dominant in direct RNA. The fact that more of these are detected at the 5’ end may be a result of the difference in coverage over transcript length between these protocols previously reported by others (41,52), which possibly leads to variation in transcript identification between the protocols. Cases of inserted exons and linked insertions are also more prevalent at the 5’ end than the 3’ end. These are cases in which the dominant transcript in the PCR cDNA has one or more exons compared with the dominant transcript in the direct RNA. Since these cases include additional splice junctions in the test transcript (dominant in PCRcDNA) they are labelled as novel by SQANTI3 (NIC or NNC). Within the transcript body (away from transcript ends), inserted or deleted exons are the most common events detected. Cases of intron retention and insertion, as well as linked deletions, insertions or combinations are also detected across all cell lines.

Finally, we want to determine whether the observed differences in transcript structure are related to the change in protocol, or if such changes are common when comparing any pair of transcripts from the same gene. For this purpose, we quantify the frequencies of structural differences detected over all the isoform switching genes in each of the cell lines, and compare them to the frequencies of structural differences detected in background genes. In order to control for the difference in mean expression between the isoform switching and background genes, for each cell line, we randomly select five sets of 100 background genes with an expression distribution similar to that of the isoform switching genes from the same cell line.

Fig. 3h shows that at the 3’ and 5’ transcript ends, isoform switching genes have fewer exon truncation events, and more exon extension in the PCR cDNA compared to background genes. These events at the transcripts’ edge are likely due to different rates of read truncation between the protocols, possibly driven by RNA degradation or PCR amplification bias. The direction of these differences does not necessarily indicate that the dominant transcript in direct RNA is always longer than the dominant transcript in PCR cDNA. While perhaps surprising, this result agrees with our findings regarding differences in transcript length mentioned earlier (Supplementary Fig. S3d), and may be due to misallocation of reads to transcript counts as shown in Supplementary Fig. S4. In the internal transcript regions not adjacent to the transcript edge, significantly more events of intron insertion and exon truncation at the 3’ end of exons are detected in the DTU pairs than in the background pairs. On the other hand, there are significantly fewer events of entire exon skipping or exon insertions in the isoform switching genes than in the background genes. It is unclear what processes in the protocols are responsible for these results and it is possible that they arise due to reverse transcription errors. In addition, some of these observations could be the result of misallocation of reads to transcript counts during the annotation/quantification process. In any case, these results highlight that, even with long read sequencing, there is still ambiguity in fully defining transcript structures.

### Transcripts in cDNA that are not observed in direct RNA are detected

In this section we analyse a much smaller group of DTU transcripts which have effectively zero proportion in direct RNA samples (prop<0.001), but are significantly higher in PCR cDNA samples from the same cell line (prop > 0.01, adjPval < 0.05). These transcripts may indicate the mechanisms that result in identification of novel transcripts in the PCR cDNA samples.

One example of a transcript that satisfies these conditions in the A549 cell line is shown in Fig. 4a (ENST00000615544). The proportion of this transcript is estimated to be effectively zero in the direct RNA, and around 0.1 in the PCR cDNA. We define this as a ‘novel isoform’, and compare it to the dominant transcript in direct RNA (ENST00000331925). Here the novel isoform is defined as the test, and the dominant transcript in direct RNA is defined as the reference. Using the same approach discussed earlier, we identify and label all structural differences found between these transcript pairs (Fig. 4c). In this example, the inserted intron, which was detected based on transcript annotation and estimated proportion, is validated qualitatively in the coverage plots.

**Figure 4:**
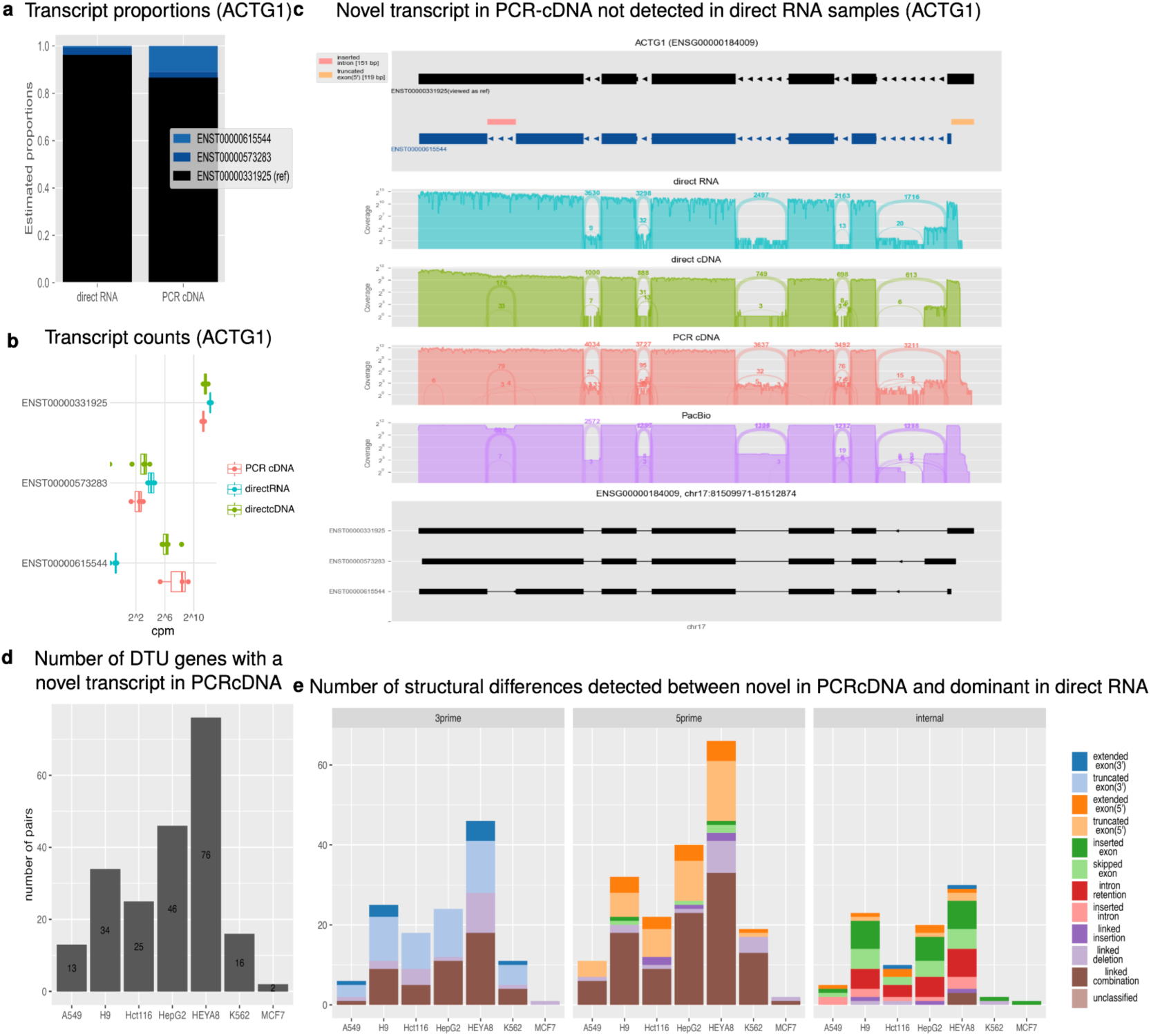
(‘Novel’ transcript only observed in PCR cDNA are structurally different to the most dominant transcript in direct RNA) a-c) Example of a DTU gene (ACTG1), where a ‘novel’ transcript was detected in PCR cDNA. a) Estimated proportions in direct RNA (left) and PCR cDNA (right), estimated using DRIMSeq from ‘bambu’ raw transcript counts for the same gene. b) Transcript counts (log2cpm) obtained by ‘bambu’ for this gene. c) When compared to the dominant transcript in direct RNA (black), the novel transcript detected in PCR cDNA (dark blue) has a 5’ truncated exon (can be seen on the right), and an inserted intron on the left. In this case, coverage plots corroborate this junction in all RT protocols, but not in direct RNA. d) Fraction of DTU genes detected, with the fraction of genes with novel isoforms in black. e) Number of structural differences detected between transcript novel in cDNA and the one dominant in RNA.

The junction defining the inserted intron in this transcript appears in all protocols that include RT, including direct cDNA and PacBio, but is absent in the direct RNA. Indeed some of the inserted intron events we observe in novel isoforms are similar to the false exitrons previously reported in (14). However, similar to in the case of isoform switching, also here, some of the novel isoforms we discover based on transcript quantification and annotation cannot be validated qualitatively in coverage plots (Supplementary Fig. S4a-c).

The fraction of genes in which such novel transcripts are identified is much lower than that of switching isoforms discussed earlier, 59/6177 (Fig. 4d). The quantification of all structural differences of these novel isoforms detected across cell lines is presented in Fig. 4e. Interestingly, while event frequencies at transcript edges seem similar to those found in the switching isoforms comparison, the frequencies of internal events detected are different. In the novel isoforms, inserted and retained introns are more common, whereas in the switching isoforms skipped and inserted exons are the most prevalent.

## Discussion

Oxford nanopore technology (ONT) can sequence full length transcripts with a variety of protocols. In addition to sequencing PCR amplified cDNA like previous technologies, ONT can also sequence RNA directly without reverse transcription or amplification. It has been shown that PCR amplification (6,7), RT (14,15), and RNA degradation (17,18) can introduce biases in RNA sequencing data, but a thorough quantification and description of their impact on transcript counts and annotations in the context of long-read transcriptome sequencing is still lacking. We perform a comprehensive comparison of transcriptomes generated with ONT sequencing using different preparation protocols, and show statistically significant differences between them in measured transcript abundance and structure.

Hundreds of genes and transcripts are found to be significantly differentially expressed between ONT sequencing protocols across seven different cell lines. Analysis of these significant DE transcripts between direct RNA and PCR cDNA showed known biases associated with expression level, sequence length and GC content. Notably, we further discovered that a higher fraction of RNA (m^6^A) methylated sites is associated with lower expression in cDNA samples.

Significant differences in transcript usage between protocols are also detected, with hundreds of DTU genes found across cell lines. However, the factors driving the bias in expression seem to play a lesser role in driving DTU. Our investigation into the structural differences between transcripts that switch dominance between the direct RNA and PCR cDNA samples reveals that while some switching events relate to simple fragmentation or truncation of the RNA in one of the protocols, most involve complex changes to the inferred structure of isoforms such as novel junctions, exon truncation, exon skipping and intron retention. The distribution of event types is different to that detected between background (non-DTU) transcripts, with fewer cases of skipped and truncated exons, but more events of intron retention and insertion (exitron). When comparing the structure of transcripts detected only in the PCR cDNA samples (i.e. novel in PCR cDNA samples) with the most dominant transcript of the same gene in the direct RNA samples, cases of inserted and retained introns are even more common. Inserted intron events (exitrons) that are only detected in PCR cDNA samples but not in their direct RNA counterparts were previously reported in (14).

The multitude of differences detected between the protocols serves to reiterate the importance of implementing the correct experimental design. Samples obtained using different sequencing protocols should not be compared with each other across different conditions, and, if they come from the same condition, should not be considered as replicates. Differential expression analysis implementing the wrong experimental design could give rise to false positive DE and DTU results driven by the biases detected between sequencing protocols, and possibly false negative results due to dilution of the true signal.

Comparing samples generated by the same protocol should counteract most of the impact of biases affecting transcript abundance since these generally impact all samples to the same extent. Indeed, we observe a high overlap in differential expression between cell lines. However, the differences in dominant transcript structure between the protocols cannot be mitigated this way, and the use of orthogonal methods to validate the existence of specific transcripts might still be required (36,38).

Since the quantification of significant differences between protocols presented in this analysis is reliant on transcript annotation and quantification tools, it is limited by their ability to provide accurate transcript identification and counts. Despite this, we show credible cases in which the structure of the dominant transcript is different between sequencing protocols, and also cases in which a transcript not detected in the direct RNA samples ‘appears’ in the PCR cDNA samples. This work is also not designed to determine which of the protocols enables a description of the transcriptome that is closer to the truth, since orthogonal methods to verify the presence of specific transcripts in a sample, i.e. laboratory method as in (14) or combination of laboratory validation with computational methods in (2), were not implemented.

Finally, our in-depth investigation of this dataset demonstrates that despite great progress in transcriptome reconstruction and quantification tools, misallocation of reads to transcripts still occurs, and leads to inaccurate transcript identification and quantification. These errors have a flow on effect to downstream analysis, leading to false identification of DTE and DTU. Therefore, with the current state and maturity of transcriptome reconstruction and quantification tools, we strongly recommend visualising read coverage data to provide a qualitative validation of events that are detected based on counts and annotation data alone.

## Conclusions

In this study, we performed a comprehensive comparison between the direct RNA and PCR-amplified cDNA ONT library preparation protocols across seven cell lines, demonstrating that protocol choice introduces substantial biases in both transcript abundance and structure. We confirmed known PCR-associated biases related to GC content and transcript length, and further identified m^6^A RNA methylation as a novel biasing factor, with highly methylated transcripts consistently down-regulated following reverse transcription and amplification. Widespread differential transcript usage was detected across all cell lines, with approximately one third of DTU genes displaying isoform switching between protocols. Structural characterisation of these switching isoforms revealed predominantly complex splicing differences rather than simple truncation, including elevated rates of intron insertion (exitron) events in PCR cDNA samples. These findings underscore the importance of consistent protocol use within experiments, the risks of misallocation of reads to transcripts affecting downstream analyses, and the value of visualising raw read coverage data as qualitative validation of annotation- and count-based results.

Taken together, these findings suggest that large-scale transcriptomic reference datasets, such as those used to define reference transcriptomes or train isoform quantification tools, may themselves carry protocol-specific biases that could propagate into downstream analyses and biological interpretations. As the field moves towards more comprehensive and accurate transcriptome atlases, careful consideration of the preparation protocol used to generate reference data will be essential to ensure that observed isoform structures and abundances reflect true biology rather than technical artefact.

## Data availability

Detailed code for the analysis and figures presented in this report is provided at https://zenodo.org/records/21132314. The data analysed here, first reported in (39) is publicly available from https://registry.opendata.aws/sg-nex-data.

## Supplementary data statement

‘Supplementary Data are available at NAR Genomics and Bioinformatics Online’.

## Acknowledgements

We would like to thank Nadia Davidson, Jonathan Göke, Jovana Maksimovic and George Howitt for helpful discussions, insights and input that helped us improve this work.

## Authors’ contributions

A.O. conceived this study, acquired funding for its execution, and provided supervision. R.A. performed research investigation, formal analysis, development of methodology, computational analyses, software development, visualisation, validation, and writing of the original draft. Both authors contributed to conceptualisation of this research, reviewing and editing of the final manuscript.

