## Supplementary material: Table 1, Supplementary Figure S1-S4 for "Long read sequencing reveals changes in measured isoform structure and abundance due to RNA processing"

Table 1: Sample inclusion table

| | library_size | protocol | grouped_protocol | cell_line | in_1a_b | in_1c_h | in_sup1a_b | in_sup1c_h | in_DE_fig2 | in_DTU_fig3 |
|---|---|---|---|---|---|---|---|---|---|---|
| SGNex_A549_cDNA_replicate1_run2 | 2290358 | cDNA | PCRcDNA | A549 | Yes | Yes | Yes | Yes | Yes | Yes |
| SGNex_A549_cDNAStranded_replicate5_run2 | 4967561 | cDNAStranded | PCRcDNA | A549 | Yes | Yes | Yes | Yes | Yes | Yes |
| SGNex_A549_directcDNA_replicate1_run3 | 1994284 | directcDNA | directcDNA | A549 | Yes | Yes | Yes | Yes | Yes | No |
| SGNex_A549_directcDNA_replicate2_run1 | 211602 | directcDNA | directcDNA | A549 | Yes | No | Yes | No | No | No |
| SGNex_A549_directcDNA_replicate3_run1 | 73369 | directcDNA | directcDNA | A549 | Yes | No | Yes | No | No | No |
| SGNex_A549_directcDNA_replicate3_run2 | 1722479 | directcDNA | directcDNA | A549 | Yes | Yes | Yes | Yes | Yes | No |
| SGNex_A549_directcDNA_replicate5_run3 | 1066082 | directcDNA | directcDNA | A549 | Yes | Yes | Yes | Yes | Yes | No |
| SGNex_A549_directRNA_replicate1_run1 | 435594 | directRNA | directRNA | A549 | Yes | Yes | Yes | Yes | Yes | Yes |
| SGNex_A549_directRNA_replicate4_run1 | 3804854 | directRNA | directRNA | A549 | Yes | Yes | Yes | Yes | Yes | Yes |
| SGNex_A549_directRNA_replicate5_run1 | 1634360 | directRNA | directRNA | A549 | Yes | Yes | Yes | Yes | Yes | Yes |
| SGNex_A549_directRNA_replicate6_run1 | 1221229 | directRNA | directRNA | A549 | Yes | Yes | Yes | Yes | Yes | Yes |
| SGNex_A549_PacBio-SMRTcell_replicate7_run1 | 1773362 | PacBio-SMRTcell | PacBio-SMRTcell | A549 | Yes | Yes | No | No | No | No |
| SGNex_H9_cDNA_replicate2_run4 | 19365758 | cDNA | PCRcDNA | H9 | Yes | Yes | Yes | Yes | Yes | Yes |
| SGNex_H9_cDNA_replicate3_run4 | 20064011 | cDNA | PCRcDNA | H9 | Yes | Yes | Yes | Yes | Yes | Yes |
| SGNex_H9_cDNA_replicate4_run4 | 18049924 | cDNA | PCRcDNA | H9 | Yes | Yes | Yes | Yes | Yes | Yes |
| SGNex_H9_directcDNA_replicate1_run2 | 614834 | directcDNA | directcDNA | H9 | Yes | Yes | Yes | Yes | Yes | No |
| SGNex_H9_directcDNA_replicate2_run3 | 12171856 | directcDNA | directcDNA | H9 | Yes | Yes | Yes | Yes | Yes | No |
| SGNex_H9_directcDNA_replicate3_run3 | 10167526 | directcDNA | directcDNA | H9 | Yes | Yes | Yes | Yes | Yes | No |
| SGNex_H9_directcDNA_replicate4_run3 | 10795954 | directcDNA | directcDNA | H9 | Yes | Yes | Yes | Yes | Yes | No |
| SGNex_H9_directRNA_replicate1_run1 | 283933 | directRNA | directRNA | H9 | Yes | No | Yes | No | No | No |
| SGNex_H9_directRNA_replicate2_run1 | 573059 | directRNA | directRNA | H9 | Yes | Yes | Yes | Yes | Yes | Yes |
| SGNex_H9_directRNA_replicate2_run2 | 280559 | directRNA | directRNA | H9 | Yes | No | Yes | No | No | No |
| SGNex_H9_directRNA_replicate3_run1 | 553044 | directRNA | directRNA | H9 | Yes | Yes | Yes | Yes | Yes | Yes |
| SGNex_H9_directRNA_replicate3_run2 | 263186 | directRNA | directRNA | H9 | Yes | No | Yes | No | No | No |
| SGNex_H9_directRNA_replicate4_run1 | 796681 | directRNA | directRNA | H9 | Yes | Yes | Yes | Yes | Yes | Yes |
| SGNex_H9_directRNA_replicate4_run2 | 188023 | directRNA | directRNA | H9 | Yes | No | Yes | No | No | No |
| SGNex_Hct116_cDNA_replicate1_run6 | 821285 | cDNA | PCRcDNA | Hct116 | Yes | Yes | Yes | Yes | Yes | Yes |
| SGNex_Hct116_cDNA_replicate3_run3 | 3608620 | cDNA | PCRcDNA | Hct116 | Yes | Yes | Yes | Yes | Yes | Yes |
| SGNex_Hct116_cDNA_replicate3_run5 | 89119957 | cDNA | PCRcDNA | Hct116 | Yes | No | Yes | No | No | No |
| SGNex_Hct116_cDNA_replicate4_run2 | 3686874 | cDNA | PCRcDNA | Hct116 | Yes | Yes | Yes | Yes | Yes | Yes |
| SGNex_Hct116_cDNA_replicate4_run4 | 79862063 | cDNA | PCRcDNA | Hct116 | Yes | No | Yes | No | No | No |
| SGNex_Hct116_cDNA_replicate4_run5 | 11701003 | cDNA | PCRcDNA | Hct116 | Yes | Yes | Yes | Yes | Yes | Yes |
| SGNex_Hct116_directcDNA_replicate1_run4 | 116151 | directcDNA | directcDNA | Hct116 | Yes | No | Yes | No | No | No |
| SGNex_Hct116_directcDNA_replicate1_run5 | 356184 | directcDNA | directcDNA | Hct116 | Yes | No | Yes | No | No | No |
| SGNex_Hct116_directcDNA_replicate3_run2 | 6184262 | directcDNA | directcDNA | Hct116 | Yes | Yes | Yes | Yes | Yes | No |
| SGNex_Hct116_directcDNA_replicate4_run1 | 6598051 | directcDNA | directcDNA | Hct116 | Yes | Yes | Yes | Yes | Yes | No |
| SGNex_Hct116_directcDNA_replicate5_run1 | 6053953 | directcDNA | directcDNA | Hct116 | Yes | Yes | Yes | Yes | Yes | No |
| SGNex_Hct116_directRNA_replicate1_run1 | 57940 | directRNA | directRNA | Hct116 | Yes | No | Yes | No | No | No |
| SGNex_Hct116_directRNA_replicate1_run2 | 520371 | directRNA | directRNA | Hct116 | Yes | Yes | Yes | Yes | Yes | Yes |
| SGNex_Hct116_directRNA_replicate1_run3 | 880168 | directRNA | directRNA | Hct116 | Yes | Yes | Yes | Yes | Yes | Yes |
| SGNex_Hct116_directRNA_replicate2_run1 | 56678 | directRNA | directRNA | Hct116 | Yes | No | Yes | No | No | No |
| SGNex_Hct116_directRNA_replicate2_run2 | 6672 | directRNA | directRNA | Hct116 | Yes | No | Yes | No | No | No |
| SGNex_Hct116_directRNA_replicate2_run3 | 173622 | directRNA | directRNA | Hct116 | Yes | No | Yes | No | No | No |
| SGNex_Hct116_directRNA_replicate2_run4 | 134350 | directRNA | directRNA | Hct116 | Yes | No | Yes | No | No | No |
| SGNex_Hct116_directRNA_replicate2_run5 | 141055 | directRNA | directRNA | Hct116 | Yes | No | Yes | No | No | No |
| SGNex_Hct116_directRNA_replicate2_run6 | 138923 | directRNA | directRNA | Hct116 | Yes | No | Yes | No | No | No |
| SGNex_Hct116_directRNA_replicate3_run1 | 1029201 | directRNA | directRNA | Hct116 | Yes | Yes | Yes | Yes | Yes | Yes |
| SGNex_Hct116_directRNA_replicate3_run4 | 981271 | directRNA | directRNA | Hct116 | Yes | Yes | Yes | Yes | Yes | Yes |
| SGNex_Hct116_directRNA_replicate4_run3 | 1605305 | directRNA | directRNA | Hct116 | Yes | Yes | Yes | Yes | Yes | Yes |
| SGNex_Hct116_directRNA_replicate6_run1 | 41 | directRNA | directRNA | Hct116 | Yes | No | Yes | No | No | No |
| SGNex_Hct116_PacBio-SMRTcell_replicate7_run1 | 1297663 | PacBio-SMRTcell | PacBio-SMRTcell | Hct116 | Yes | Yes | No | No | No | No |
| SGNex_HepG2_cDNA_replicate1_run4 | 2481367 | cDNA | PCRcDNA | HepG2 | Yes | Yes | Yes | Yes | Yes | Yes |
| SGNex_HepG2_cDNAStranded_replicate4_run3 | 820879 | cDNAStranded | PCRcDNA | HepG2 | Yes | Yes | Yes | Yes | Yes | Yes |
| SGNex_HepG2_cDNAStranded_replicate5_run4 | 5067753 | cDNAStranded | PCRcDNA | HepG2 | Yes | Yes | Yes | Yes | Yes | Yes |
| SGNex_HepG2_directcDNA_replicate1_run1 | 507295 | directcDNA | directcDNA | HepG2 | Yes | Yes | Yes | Yes | Yes | No |
| SGNex_HepG2_directcDNA_replicate1_run2 | 983476 | directcDNA | directcDNA | HepG2 | Yes | Yes | Yes | Yes | Yes | No |

Table 1: Sample inclusion table *(continued)*

| | library_size | protocol | grouped_protocol | cell_line | in_1a_b | in_1c_h | in_sup1a_b | in_sup1c_h | in_DE_fig2 | in_DTU_fig3 |
|---|---|---|---|---|---|---|---|---|---|---|
| SGNex_HepG2_directcDNA_replicate4_run1 | 143232 | directcDNA | directcDNA | HepG2 | Yes | No | Yes | No | No | No |
| SGNex_HepG2_directcDNA_replicate4_run2 | 534401 | directcDNA | directcDNA | HepG2 | Yes | Yes | Yes | Yes | Yes | No |
| SGNex_HepG2_directcDNA_replicate5_run3 | 1295543 | directcDNA | directcDNA | HepG2 | Yes | Yes | Yes | Yes | Yes | No |
| SGNex_HepG2_directRNA_replicate1_run3 | 1108309 | directRNA | directRNA | HepG2 | Yes | Yes | Yes | Yes | Yes | Yes |
| SGNex_HepG2_directRNA_replicate2_run1 | 164770 | directRNA | directRNA | HepG2 | Yes | No | Yes | No | No | No |
| SGNex_HepG2_directRNA_replicate3_run1 | 115650 | directRNA | directRNA | HepG2 | Yes | No | Yes | No | No | No |
| SGNex_HepG2_directRNA_replicate5_run1 | 5972252 | directRNA | directRNA | HepG2 | Yes | Yes | Yes | Yes | Yes | Yes |
| SGNex_HepG2_directRNA_replicate5_run2 | 1771190 | directRNA | directRNA | HepG2 | Yes | Yes | Yes | Yes | Yes | Yes |
| SGNex_HepG2_directRNA_replicate6_run1 | 1396710 | directRNA | directRNA | HepG2 | Yes | Yes | Yes | Yes | Yes | Yes |
| SGNex_HepG2_PacBio-SMRTcell_replicate7_run1 | 2319306 | PacBio-SMRTcell | PacBio-SMRTcell | HepG2 | Yes | Yes | No | No | No | No |
| SGNex_HEYA8_cDNA_replicate1_run4 | 21843017 | cDNA | PCRcDNA | HEYA8 | Yes | Yes | Yes | Yes | Yes | Yes |
| SGNex_HEYA8_cDNA_replicate2_run4 | 21475884 | cDNA | PCRcDNA | HEYA8 | Yes | Yes | Yes | Yes | Yes | Yes |
| SGNex_HEYA8_cDNA_replicate3_run3 | 19579635 | cDNA | PCRcDNA | HEYA8 | Yes | Yes | Yes | Yes | Yes | Yes |
| SGNex_HEYA8_directcDNA_replicate1_run3 | 9162620 | directcDNA | directcDNA | HEYA8 | Yes | Yes | Yes | Yes | Yes | No |
| SGNex_HEYA8_directcDNA_replicate2_run3 | 12763098 | directcDNA | directcDNA | HEYA8 | Yes | Yes | Yes | Yes | Yes | No |
| SGNex_HEYA8_directcDNA_replicate3_run2 | 8598703 | directcDNA | directcDNA | HEYA8 | Yes | Yes | Yes | Yes | Yes | No |
| SGNex_HEYA8_directRNA_replicate1_run1 | 968474 | directRNA | directRNA | HEYA8 | Yes | Yes | Yes | Yes | Yes | Yes |
| SGNex_HEYA8_directRNA_replicate1_run2 | 236002 | directRNA | directRNA | HEYA8 | Yes | No | Yes | No | No | No |
| SGNex_HEYA8_directRNA_replicate2_run1 | 873095 | directRNA | directRNA | HEYA8 | Yes | Yes | Yes | Yes | Yes | Yes |
| SGNex_HEYA8_directRNA_replicate2_run2 | 531611 | directRNA | directRNA | HEYA8 | Yes | Yes | Yes | Yes | Yes | Yes |
| SGNex_HEYA8_directRNA_replicate3_run1 | 1207796 | directRNA | directRNA | HEYA8 | Yes | Yes | Yes | Yes | Yes | Yes |
| SGNex_HEYA8_PacBio-SMRTcell_replicate4_run1 | 2553104 | PacBio-SMRTcell | PacBio-SMRTcell | HEYA8 | Yes | Yes | No | No | No | No |
| SGNex_K562_cDNA_replicate1_run3 | 1017881 | cDNA | PCRcDNA | K562 | Yes | Yes | Yes | Yes | Yes | Yes |
| SGNex_K562_cDNA_replicate3_run4 | 6354389 | cDNA | PCRcDNA | K562 | Yes | Yes | Yes | Yes | Yes | Yes |
| SGNex_K562_cDNAStranded_replicate2_run3 | 1105400 | cDNAStranded | PCRcDNA | K562 | Yes | Yes | Yes | Yes | Yes | Yes |
| SGNex_K562_cDNAStranded_replicate3_run2 | 874321 | cDNAStranded | PCRcDNA | K562 | Yes | No | Yes | No | No | No |
| SGNex_K562_cDNAStranded_replicate3_run3 | 758658 | cDNAStranded | PCRcDNA | K562 | Yes | No | Yes | No | No | No |
| SGNex_K562_directcDNA_replicate1_run2 | 2097500 | directcDNA | directcDNA | K562 | Yes | Yes | Yes | Yes | Yes | No |
| SGNex_K562_directcDNA_replicate2_run1 | 243444 | directcDNA | directcDNA | K562 | Yes | No | Yes | No | No | No |
| SGNex_K562_directcDNA_replicate2_run2 | 205257 | directcDNA | directcDNA | K562 | Yes | No | Yes | No | No | No |
| SGNex_K562_directcDNA_replicate3_run1 | 506756 | directcDNA | directcDNA | K562 | Yes | Yes | Yes | Yes | Yes | No |
| SGNex_K562_directcDNA_replicate4_run2 | 2024500 | directcDNA | directcDNA | K562 | Yes | Yes | Yes | Yes | Yes | No |
| SGNex_K562_directRNA_replicate1_run1 | 553040 | directRNA | directRNA | K562 | Yes | Yes | Yes | Yes | Yes | Yes |
| SGNex_K562_directRNA_replicate4_run1 | 933840 | directRNA | directRNA | K562 | Yes | Yes | Yes | Yes | Yes | Yes |
| SGNex_K562_directRNA_replicate5_run1 | 869196 | directRNA | directRNA | K562 | Yes | Yes | Yes | Yes | Yes | Yes |
| SGNex_K562_directRNA_replicate6_run1 | 1331510 | directRNA | directRNA | K562 | Yes | Yes | Yes | Yes | Yes | Yes |
| SGNex_K562_PacBio-SMRTcell_replicate7_run1 | 1252301 | PacBio-SMRTcell | PacBio-SMRTcell | K562 | Yes | Yes | No | No | No | No |
| SGNex_MCF7_cDNA_replicate1_run3 | 1916066 | cDNA | PCRcDNA | MCF7 | Yes | Yes | Yes | Yes | Yes | Yes |
| SGNex_MCF7_cDNAStranded_replicate2_run1 | 1326211 | cDNAStranded | PCRcDNA | MCF7 | Yes | Yes | Yes | Yes | Yes | Yes |
| SGNex_MCF7_cDNAStranded_replicate3_run2 | 2418673 | cDNAStranded | PCRcDNA | MCF7 | Yes | Yes | Yes | Yes | Yes | Yes |
| SGNex_MCF7_directcDNA_replicate1_run2 | 847599 | directcDNA | directcDNA | MCF7 | Yes | Yes | Yes | Yes | Yes | No |
| SGNex_MCF7_directcDNA_replicate3_run3 | 2084399 | directcDNA | directcDNA | MCF7 | Yes | Yes | Yes | Yes | Yes | No |
| SGNex_MCF7_directcDNA_replicate4_run2 | 4575376 | directcDNA | directcDNA | MCF7 | Yes | Yes | Yes | Yes | Yes | No |
| SGNex_MCF7_directRNA_replicate1_run1 | 23534 | directRNA | directRNA | MCF7 | Yes | No | Yes | No | No | No |
| SGNex_MCF7_directRNA_replicate2_run2 | 271235 | directRNA | directRNA | MCF7 | Yes | No | Yes | No | No | No |
| SGNex_MCF7_directRNA_replicate2_run3 | 5014547 | directRNA | directRNA | MCF7 | Yes | Yes | Yes | Yes | Yes | Yes |
| SGNex_MCF7_directRNA_replicate3_run1 | 1350547 | directRNA | directRNA | MCF7 | Yes | Yes | Yes | Yes | Yes | No |
| SGNex_MCF7_directRNA_replicate4_run1 | 1148896 | directRNA | directRNA | MCF7 | Yes | Yes | Yes | Yes | Yes | No |
| SGNex_MCF7_PacBio-SMRTcell_replicate5_run1 | 1380802 | PacBio-SMRTcell | PacBio-SMRTcell | MCF7 | Yes | Yes | No | No | No | No |
| SGNex_MCF7-EV_directRNA_replicate1_run1 | 1923485 | directRNA | directRNA | MCF7-EV | No | No | No | No | No | No |
| SGNex_MCF7-EV_directRNA_replicate2_run1 | 1448947 | directRNA | directRNA | MCF7-EV | No | No | No | No | No | No |
| Total | 0 | NA | NA | NA | 105 | 77 | 99 | 71 | 71 | 46 |

### Supplementary Figures

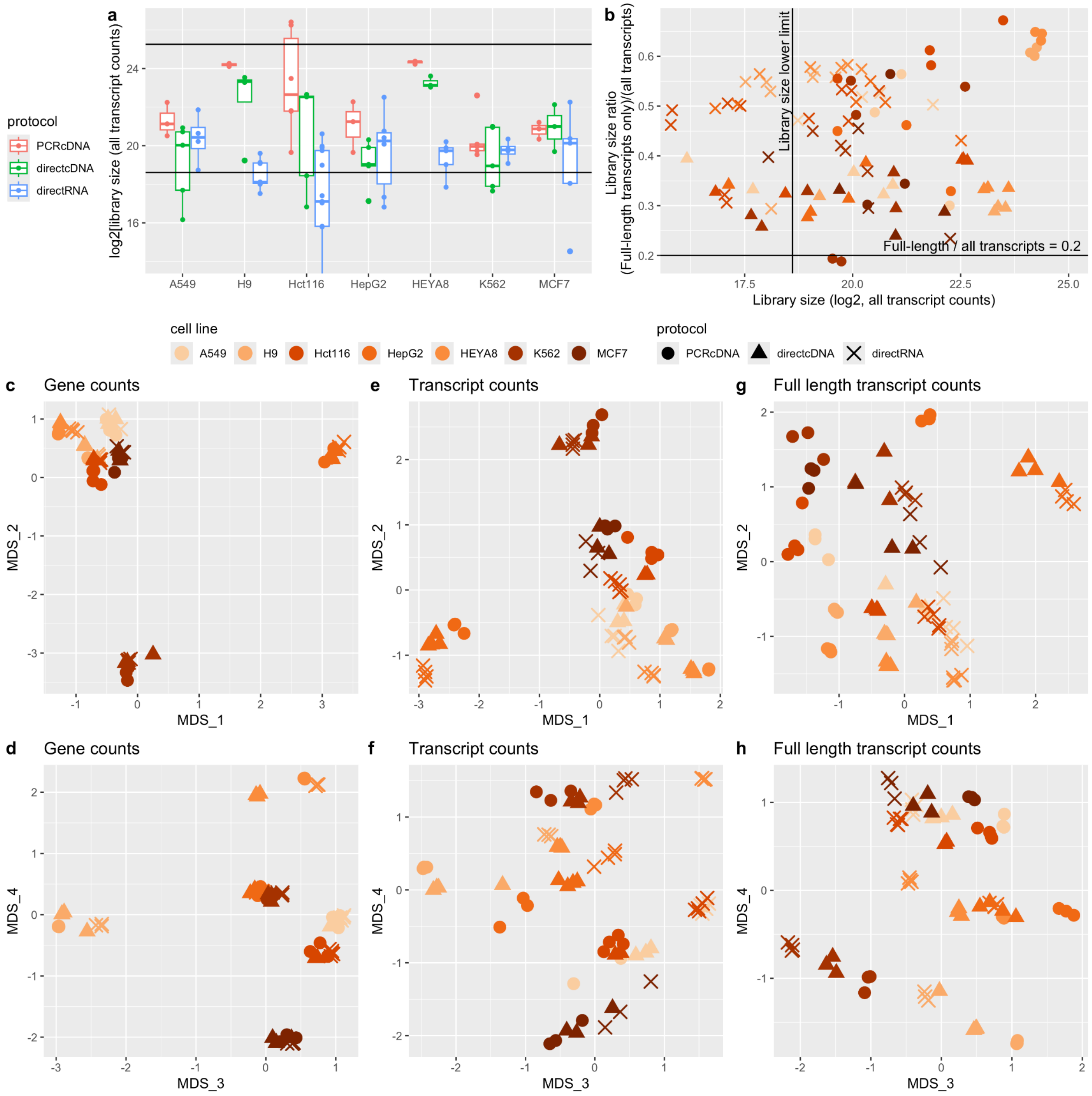


### Supplementary Figure S1

a) Library size (number of mapped reads) in each sample, across preparation protocols and cell lines. The samples between the horizontal lines were retained for analysis. b) Fraction of full length transcript reads as a function of log2(library size). MDS plots of gene counts (c,d), total transcript counts (e,f) and full length transcript counts (g,h). Samples separate based on cell line (colour) but also on preparation protocol (shape) especially at the isoform level.

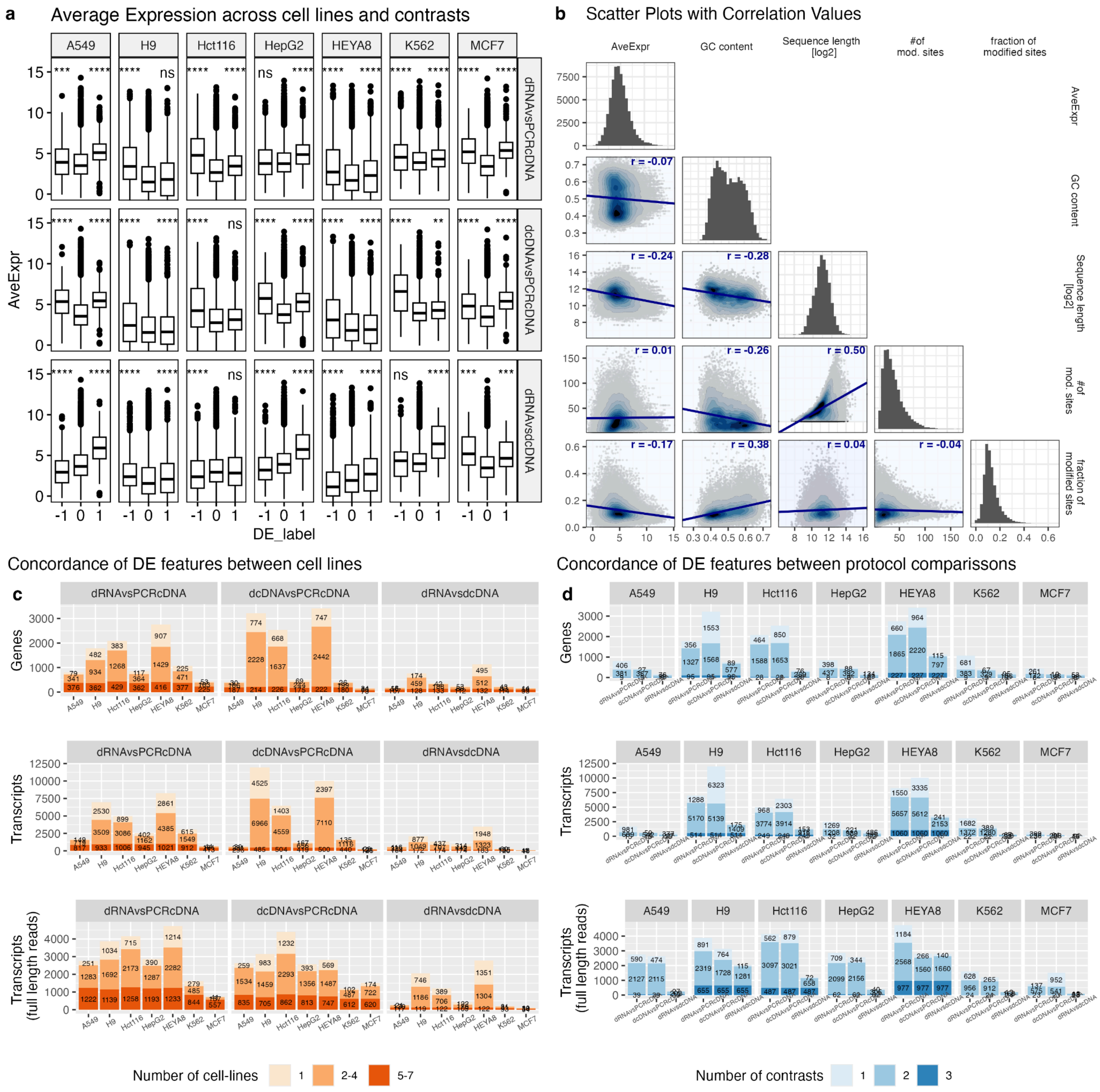


### Supplementary Figure S2

a) Mean transcript expression, stratified by DE labels across cell lines and contrasts. Significant DE features are biased towards higher expression levels as expected. P-values were obtained using a two-tailed t.test, with respect to the non DE group. ns: p > 0.05, *: p <= 0.05, **: p <= 0.01, ***: p <= 0.001, and ****: p <= 0.0001. b) Cross correlation plots of observed transcript expression, GC content, transcript length and m6A methylation. Observed average expression shows negative correlation with sequence length (r=-.24) which is expected in long read data. The number of m6A modified sites increases with transcript length (r=0.5), and decreases with GC content (r=-0.26). The fraction of m6A modified sites (which is the measure of m6A modification plotted in Fig. 2f) is not correlated with sequence length (r=0.04), and has positive correlation with GC content (r = 0.38). Average expression shows negative correlation with the fraction of modified sites (r = -0.17), but is not correlated with the number of modified sites (r=0.01). Correlation is estimated using the Pearson method. c) Concordance of DE features between cell lines. Top panel: Gene counts, middle panel: Transcript counts, bottom panel: Full length transcript counts). For each of the contrasts, the coloured bars indicate for each feature, in how many cell lines it was found to be differentially expressed between the protocols compared. One indicates the feature was found differentially expressed between the protocols compared only in that cell line (low concordance), while 5-7 indicates the feature was found differentially expressed between these protocols in all or almost all the cell lines (high concordance). For example, in the direct RNA vs PCR cDNA comparison, there are about 400 genes and 1000 transcripts that are consistently differentially expressed in 5-7 of the cell lines, which we consider high concordance. Further, there are more than 1000 genes and 4000 transcripts that are differentially expressed between the direct RNA and PCR cDNA protocols in 2-4 of the cell lines. This strong overlap of DE genes and transcripts between the cell lines, despite the large variation in expression between them gives a strong indication that the factors driving these differences are consistent across the cell lines. Indeed GC content and sequence length are intrinsic characteristics of each gene and transcript and do not change between the cell lines. d) Similar to c), but for each of the cell lines, the colour indicates how many contrasts each feature was DE in.

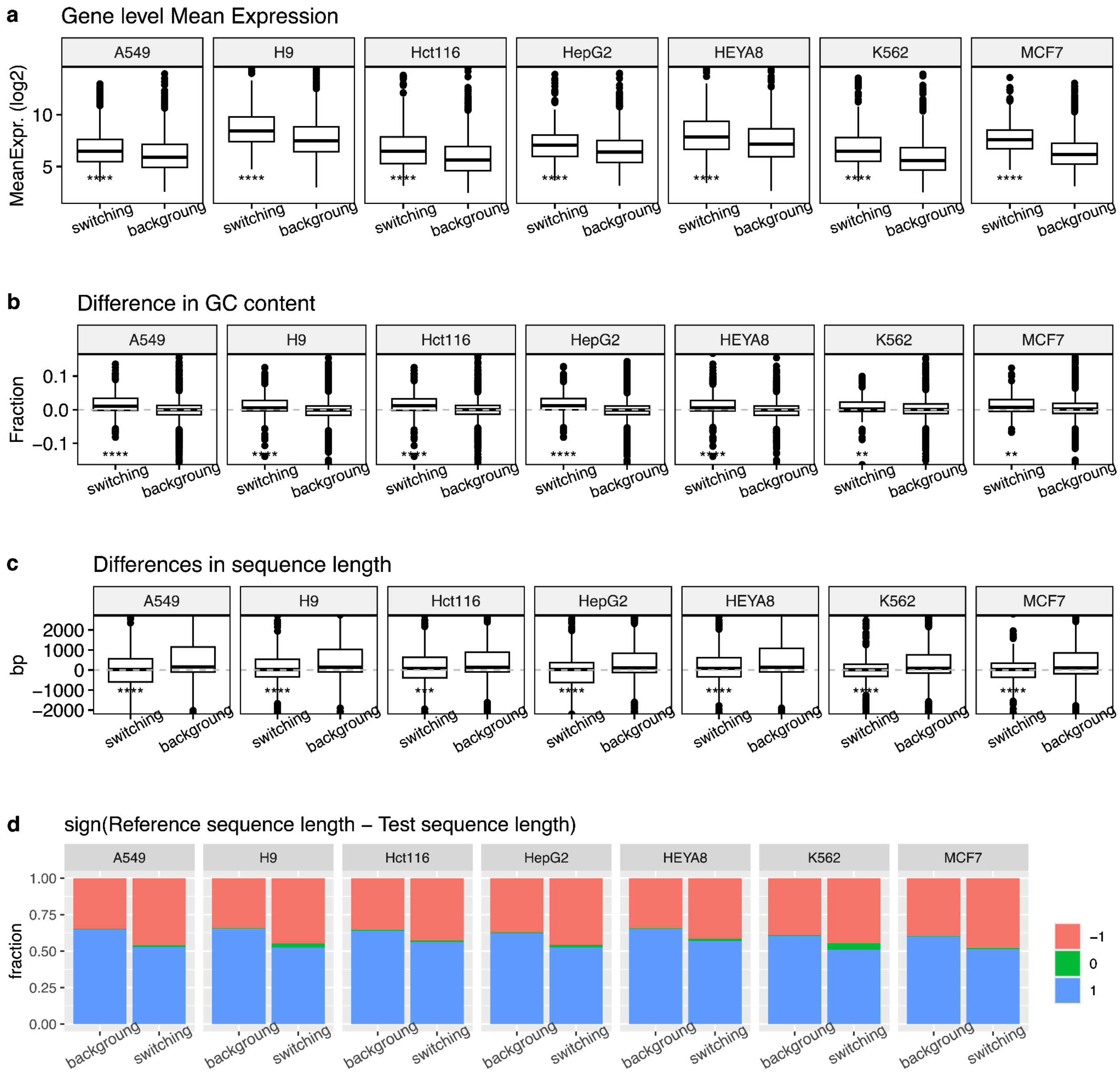


### Supplementary Figure S3

a) Gene level Mean expression split to background and dominant switching gene.
b) Difference in GC content between 'reference' and 'test' transcripts calculated for each gene, split to background and dominant switching. c) Difference in sequence length between 'reference' and 'test', in background and dominant switching genes. P-values were obtained using a two-tailed t.test, with respect to the background group. ns: p > 0.05, *: p <= 0.05, **: p <= 0.01, ***:p<= 0.001, and ****: p <= 0.0001.
d) Sign of difference in length between the 'reference' and 'test' transcripts, split to background and dominant switching. The +1 sign (blue bar) represents genes in which the 'reference' transcript (most dominant in direct RNA) is longer than the 'test' transcript (most dominant transcript in cDNA

for switching, and second most dominant in  dRNA for background transcript pairs). -1 or red bars represent the opposite case where the ‘reference’ transcript is shorter than the ‘test’ transcript. In around 60% of the background transcript pairs the transcript dominant in the direct RNA is longer. While in the switching transcript pairs it is closer to 50% for all cell types.

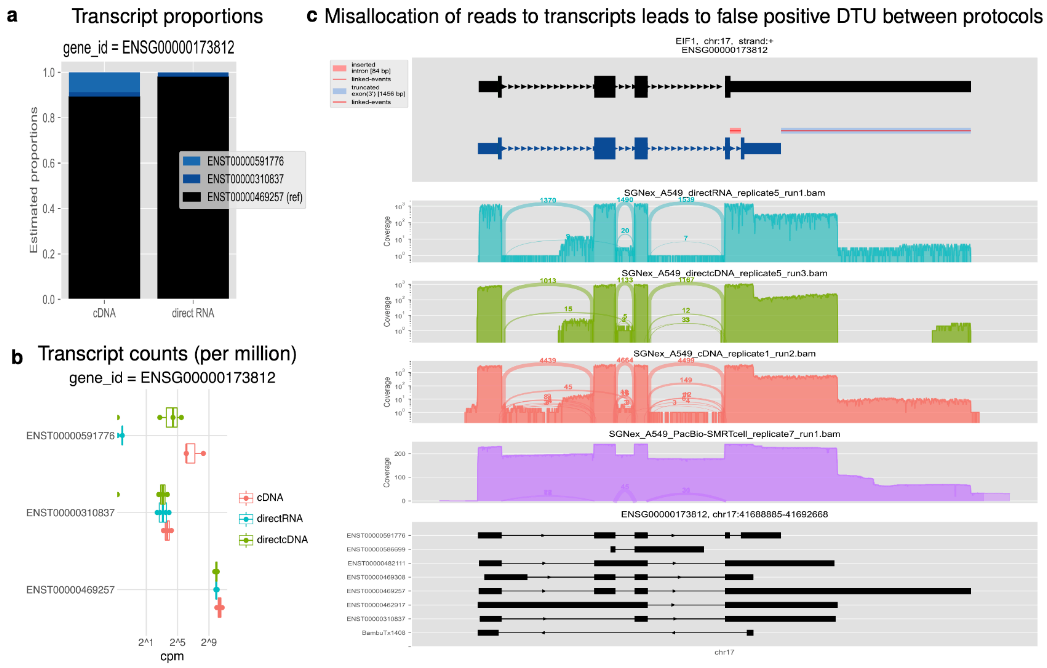


### Supplementary Figure S4

Examples of false negative detection of DTU gene due to misallocation of reads to transcripts which lead to errors in transcript quantification, these result in wrong estimation of transcript proportions, which appear to be significantly different between the protocols. a) Transcript counts b) Estimated transcript proportions c) Structural differences between dominant isoform in direct RNA and novel in PCR cDNA.
